# SAMHD1 enhances HIV-1-induced glycolysis in monocytic cells through hexokinase 2 upregulation

**DOI:** 10.64898/2026.08.03.742570

**Authors:** Hua Yang, Pak-Hin Hinson Cheung, Li Wu

## Abstract

SAMHD1 is a mitochondria-associated cellular protein that restricts HIV-1 replication by depleting intracellular dNTP pools in non-dividing immune cells, such as macrophages, dendritic cells, and resting CD4^+^ T cells; however, its role in host metabolism remains unclear. Building on our previous finding that SAMHD1 promotes mitochondrial membrane damage in HIV-1-infected monocytic cells, here we identify a new function for SAMHD1 in enhancing HIV-1-induced glycolysis through upregulation of hexokinase 2 (HK2). In monocytic THP-1 cells, but not differentiated macrophage-like cells, SAMHD1 amplifies HIV-1-triggered glucose uptake and basal glycolysis. Mechanistically, SAMHD1 increases HK2 expression and promotes its cytosolic accumulation, leading to elevated reactive oxygen species (ROS) production. This SAMHD1-dependent metabolic rewiring links antiviral restriction to glycolytic control and cellular stress responses. Our findings reveal a cell state-specific role for SAMHD1 in regulating glycolysis during HIV-1 infection, identify HK2 as a key effector, and uncover an unanticipated layer of host-virus interaction in monocytic cells.

**IMPORTANCE:** SAMHD1 is best known as a restriction factor that inhibits HIV-1 replication mainly through its dNTPase activity. However, emerging evidence suggests that SAMHD1 also regulates mitochondrial homeostasis and cellular metabolism. We previously demonstrated that SAMHD1 promotes HIV-1-induced apoptosis in monocytic cells through a mitochondrial pathway, implicating SAMHD1 in the control of mitochondrial function during infection. Because mitochondria are central regulators of cellular energy metabolism, we investigated whether SAMHD1 influences glycolytic reprogramming in HIV-1-infected monocytic cells. Our results show that SAMHD1 enhances glucose uptake, glycolysis, HK2 expression, and ROS production during HIV-1 infection. These findings reveal a previously unrecognized role for SAMHD1 in coordinating metabolic and oxidative stress responses to HIV-1 infection and provide new mechanistic insight into the interplay between antiviral factors, cellular metabolism, and HIV-1 pathogenesis. Understanding how SAMHD1 regulates glucose metabolism may uncover novel links between innate immune defenses and metabolic disease.

## INTRODUCTION

Sterile alpha motif (SAM) and histidine-aspartate (HD) domain-containing protein 1 (SAMHD1) is a dNTP triphosphohydrolase that reduces the intracellular dNTP pool,(1, 2) and restricts human immunodeficiency virus type 1 (HIV-1) in non-dividing immune cells, such as macrophages, dendritic cells, and resting CD4^+^ T cells (3–5). SAMHD1 also restricts the other retroviruses (6), including HIV-2 and certain types of simian immunodeficiency viruses (7, 8), DNA viruses, including hepatitis B virus (HBV) (9–11), human papillomavirus 16 (12), as well as RNA viruses, including influenza A virus (13, 14), hepatitis C virus and other flaviviruses (15). Conversely, SAMHD1 increases infection of severe acute respiratory syndrome coronavirus-2 in human macrophages, phorbol 12-myristate 13-acetate (PMA)-differentiated macrophage-like cells, and lung epithelial Calu-3 cells (16, 17). SAMHD1 also plays multifaceted regulatory roles in cellular processes, including innate immune responses (18–20), DNA replication fork progression (21), cell proliferation (22, 23), and glucose metabolism (24–26).

Glucose metabolism is a critical biochemical process for maintaining energy balance, which involves biochemical reactions and metabolic pathways, such as glucose uptake, glycolysis, pentose phosphate pathway, glycogenolysis, tricarboxylic acid (TCA) cycle, and oxidative phosphorylation (OXPHOS) (27–29). Glucose enters cells through glucose transporters (Gluts) and is phosphorylated to glucose-6-phosphate by hexokinase 1 and hexokinase 2 (HK1/2), which is the first step of glycolysis. Unlike HK1, which is broadly expressed in most adult tissues, HK2 exhibits a more tissue-specific expression pattern and is highly expressed in skeletal muscle cells, heart cells, adipocytes, and activated T cells (27, 30). Pyruvate, the final product of glycolysis, enters mitochondria and fuels the TCA cycle. The TCA cycle generates nicotinamide adenine dinucleotide and flavin adenine dinucleotide, which drive the synthesis of OXPHOS and adenosine triphosphate (ATP) (28, 31).

HIV-1 selectively infects metabolically active CD4^+^ T cells (32). HIV-1 infection in primary CD4^+^ T cells modulates glucose metabolism by upregulating glucose uptake and hexokinase activity (33), although the underlying molecular mechanisms remain to be determined. Mitochondria are essential organelles in glucose metabolism responsible for ATP production, reactive oxygen species (ROS) generation, and regulation of cell death (34). Our previous studies demonstrated that endogenous SAMHD1 promotes both spontaneous apoptosis and HIV-1-induced apoptosis through the mitochondrial pathway in monocytic THP-1 cells (22, 35), establishing this cell line as a suitable model for investigating the mechanisms by which SAMHD1 regulates HIV-1 infection and cellular responses in monocytic cells. Particularly, SAMHD1 is localized to the mitochondria and promotes mitochondrial membrane damage in HIV-1-infected THP-1 cells (20, 35). However, the functions and mechanisms of SAMHD1 in regulating glucose metabolism in HIV-1-infetcted cells remain unclear, which represents a significant knowledge gap in our understanding of HIV-1 modulation of cellular metabolism.

In this study, we find that endogenous SAMHD1 enhances HIV-1-induced metabolic reprogramming through the regulation of HK2 expression and subcellular localization in THP-1 cells. We demonstrate that SAMHD1 promotes HIV-1-induced glucose uptake and basal glycolysis. Furthermore, SAMHD1-enhanced glycolysis is associated with an increased HK2 expression and the accumulation of cytosolic HK2, which leads to ROS production in HIV-1-infected cells. Our findings reveal a cellular mechanism by which SAMHD1 regulates glycolysis and ROS production in HIV-1-infected monocytic cells, suggesting a previously unrecognized function of SAMHD1 during HIV-1 infection.

## RESULTS

### Endogenous SAMHD1 enhances HIV-1-induced glucose uptake in dividing THP-1 cells, but not in differentiated THP-1 cells

SAMHD1 is a cellular restriction factor against HIV-1 infection mainly in non-dividing myeloid cells (3, 4). Our previous study showed that SAMHD1 efficiently inhibits a single-cycle, luciferase reporter HIV-1 (HIV-1-Luc/VSV-G) infection in PMA-differentiated THP-1 cells, which resembles non-dividing macrophage-like cells (22). Of note, SAMHD1 inhibits early HIV-1-Luc/VSV-G infection at 1 day post-infection (dpi) in dividing THP-1 cells, but not at later time point at 2 dpi (22). Using the same luciferase reporter HIV-1 in the current study, we found that HIV-1-infected THP-1 control (Ctrl) cells exhibited lower luciferase activity than SAMHD1 knockout (KO) THP-1 cells at 1 dpi (Fig. 1A). In contrast, luciferase activities were slightly higher in Ctrl cells than in SAMHD1 KO cells at 2-4 dpi. Following PMA differentiation of cells, HIV-1 infection was markedly enhanced in SAMHD1 KO cells compared with Ctrl cells at both 1 and 2 dpi (Fig. 1B). As a negative control, treatment of the cells with the reverse transcriptase inhibitor nevirapine (NVP) completely abolished HIV-1 replication (Fig. 1A and 1B).

**Fig. 1.**
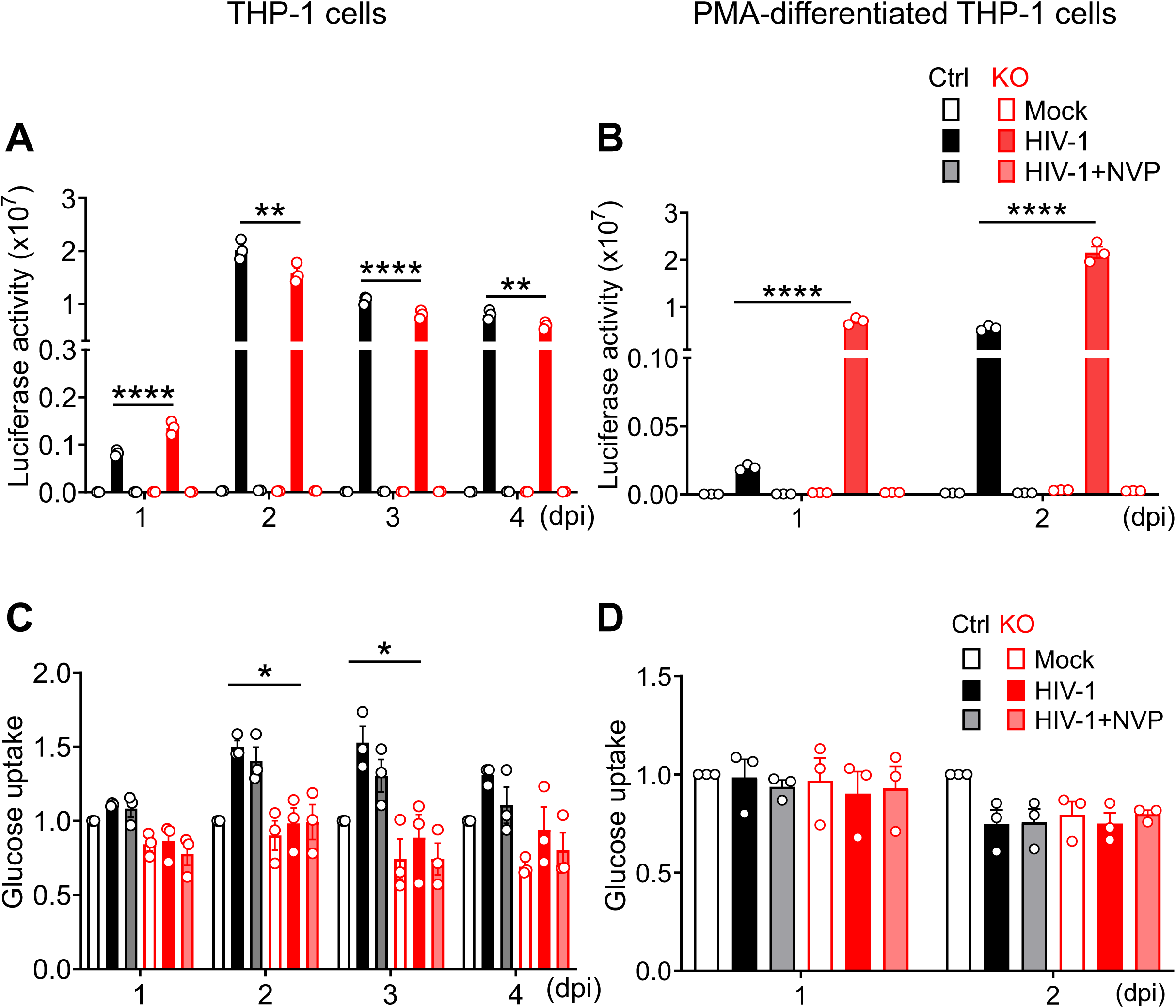
Endogenous SAMHD1 enhances HIV-1-induced glucose uptake in THP-1 cells, not in differentiated cells. (A, C) THP-1 control (Ctrl) cells and SAMHD1 knockout (KO) cells, (B, D) PMA-differentiated THP-1 Ctrl and SAMHD1 KO cells were infected with single-cycle, luciferase reporter HIV-1-Luc/VSV-G (multiplicity of infection [MOI] = 2) or mock infected. NVP was used to inhibit HIV-1 reverse transcription and infection. THP-1 cells were harvested at 1-4 dpi and differentiated THP-1 cells were harvested at 1-2 dpi for analyses. (A, B) HIV-1 infection levels were measured by the luciferase assay and normalized to the amount of cellular protein. (C, D) Glucose uptake was measured by 2-NBDG staining and flow cytometry. The level of mock in Ctrl groups was set as 1. Original flow cytometry results of glucose uptake are shown in Fig. S1. (A-D) Data are presented as means ± SEM from three independent experiments. The one-way ANOVA was used for statistical significance compared with THP-1 Ctrl cells with HIV-1 infection. \**P* < 0.05, \*\**P* < 0.01, \*\*\*\**P* < 0.0001.

HIV-1 infection of CD4^+^ T cells or macrophages promotes glucose uptake, although the underlying mechanisms remain to be defined (36–38). To investigate the effect of SAMHD1 on HIV-1-induced glucose uptake, Ctrl cells and SAMHD1 KO cells or PMA-differentiated counterpart cells were infected with HIV-1 and treated with or without NVP. Glucose uptake was measured by staining of 2-[*N*-(7-nitrobenz-2-oxa-1,3-diazol-4-yl) amino]-2-deoxy-d-glucose (2-NBDG) using flow cytometry. Interestingly, HIV-1 infection promoted glucose uptake in Ctrl cells at 2 and 3 dpi, which was mitigated in SAMHD1 KO cells (Fig. 1C and Fig. S1A), suggesting that SAMHD1 enhances HIV-1-induced glucose uptake in THP-1 cells. Of note, NVP treatment did not reduce the HIV-1-induced glucose uptake in Ctrl cells (Fig. 1C and Fig. S1A), suggesting that this SAMHD1-enhanced glucose uptake is independent of HIV-1 replication. Conversely, in PMA-differentiated cells, glucose uptake was not affected by HIV-1 infection, SAMHD1 expression, or NVP treatment (Fig. 1D and Fig. S1B). These results indicate that endogenous SAMHD1 enhances HIV-1-induced glucose uptake in dividing THP-1 cells, but not in differentiated macrophage-like cells.

### SAMHD1 enhances HIV-1-induced basal glycolysis in THP-1 cells, but not in differentiated cells

HIV-1 infection can promote glycolysis in CD4^+^ T cells (32, 37). To determine whether SAMHD1 affects glycolysis in monocytic cells upon HIV-1 infection, Ctrl cells and SAMHD1 KO cells were infected with HIV-1 and analyzed at 1 and 2 dpi. Glycolytic proton extrusion rate (GlycoPER) was measured with a Seahorse XFe24 analyzer and used to evaluate basal glycolysis that reflects glycolytic activity under basal conditions, and compensatory glycolysis that reflects forced glycolysis after mitochondrial inhibition by rotenone and antimycin A (39). We observed that HIV-1-induced basal glycolysis was stronger in Ctrl cells (a 2.4-fold increase) than in SAMHD1 KO cells (a 1.5-fold increase) at 1 dpi (Fig. 2A and 2C), despite lower HIV-1 infection in Ctrl cells than SAMHD1 KO cells (Fig. 1A), suggesting that SAMHD1 enhances HIV-1-induced basal glycolysis in THP-1 cells at 1 dpi. Conversely, HIV-1-induced compensatory glycolysis was less affected by SAMHD1 (∼1.6 fold in Ctrl vs. ∼1.3-fold in KO cells) (Fig. 2A and 2E). At 2 dpi, uninfected THP-1 cells showed increased basal and compensatory glycolysis, which could not be further induced by HIV-1 infection (Fig. 2B, 2D, and 2F). At 2 dpi, HIV-1 infection decreased the compensatory glycolysis in Ctrl cells, but not in SAMHD1 KO cells (Fig. 2B and 2F).

**Fig. 2.**
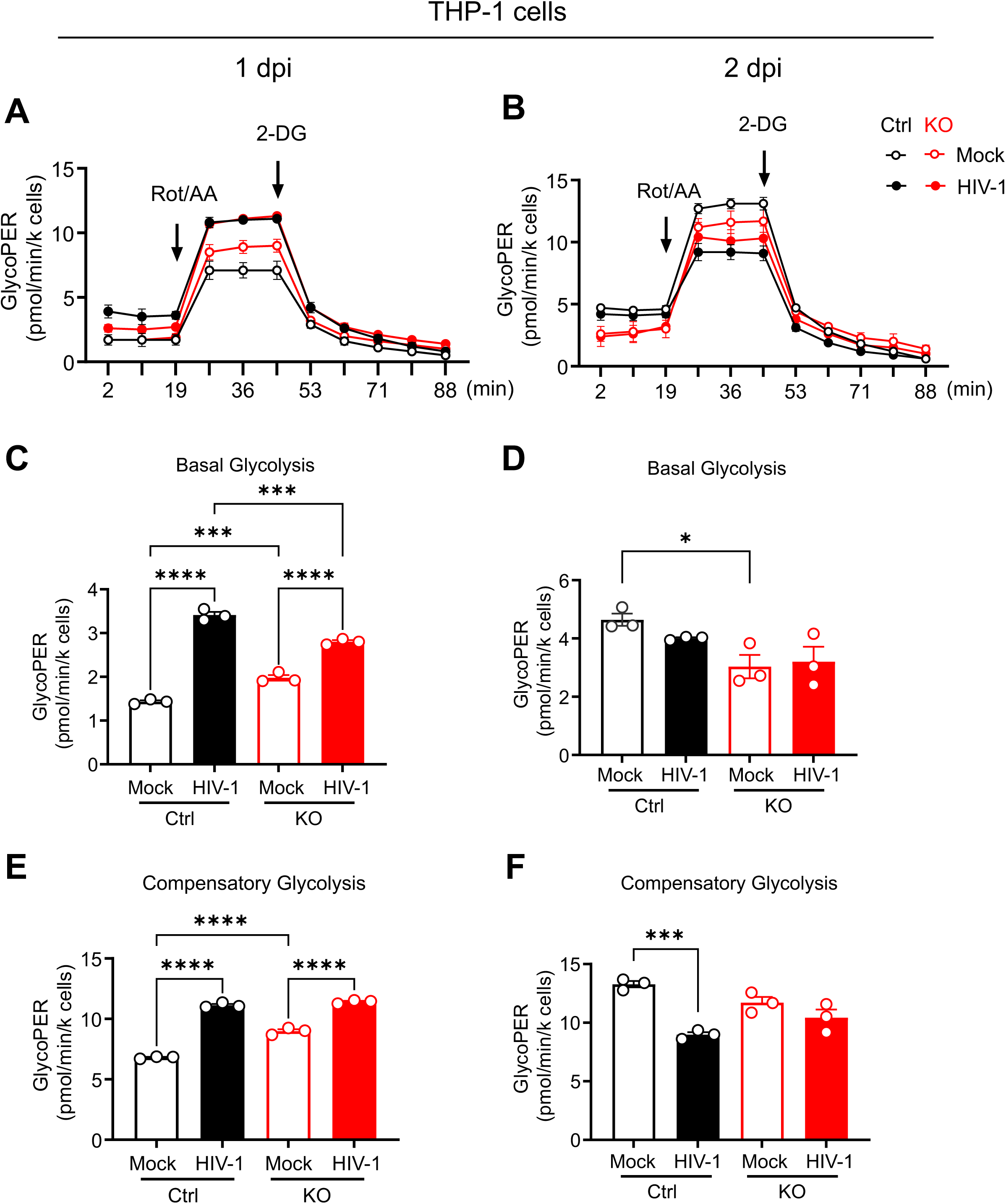
SAMHD1 enhances HIV-1-induced basal glycolysis in THP-1 cells. THP-1 Ctrl and SAMHD1 KO cells were infected with HIV-1-Luc/VSV-G (MOI = 2) or mock infected. Cells were harvested at 1 dpi (A, C, E) or 2 dpi (B, D, F) for analyses. (A, B) GlycoPER was measured by a Seahorse XFe24 analyzer at indicated time points and used to evaluate the basal and compensatory glycolysis levels. (C-F) The average values of basal glycolysis (C, D) and compensatory glycolysis (E, F) levels of THP-1 Ctrl and SAMHD1 KO cells with HIV-1 infection or mock infected. Data are presented as means ± SEM from three independent experiments. (C-F) One-way ANOVA was used for statistical significance. \**P* < 0.05, \*\*\**P* < 0.001, \*\*\*\**P* < 0.0001.

To investigate the effect of SAMHD1 on glycolysis in non-dividing macrophage-like cells, PMA-differentiated Ctrl cells and SAMHD1 KO cells were infected with HIV-1 and analyzed at 1 dpi and 2 dpi. We found that neither basal glycolysis nor compensatory glycolysis was affected by HIV-1 infection or SAMHD1 expression at 1 dpi (Fig. 3A, 3C and 3E). In contrast, at 2 dpi, both basal glycolysis and compensatory glycolysis were increased by HIV-1 infection in differentiated SAMHD1 KO cells, but not in differentiated Ctrl cells (Fig. 3B, 3D and 3F), which were consistent with higher HIV-1 infection in SAMHD1 KO cells (Fig. 1B). Together, these results suggest that endogenous SAMHD1 enhances HIV-1-induced basal glycolysis in THP-1 cells, but not in differentiated cells.

**Fig. 3.**
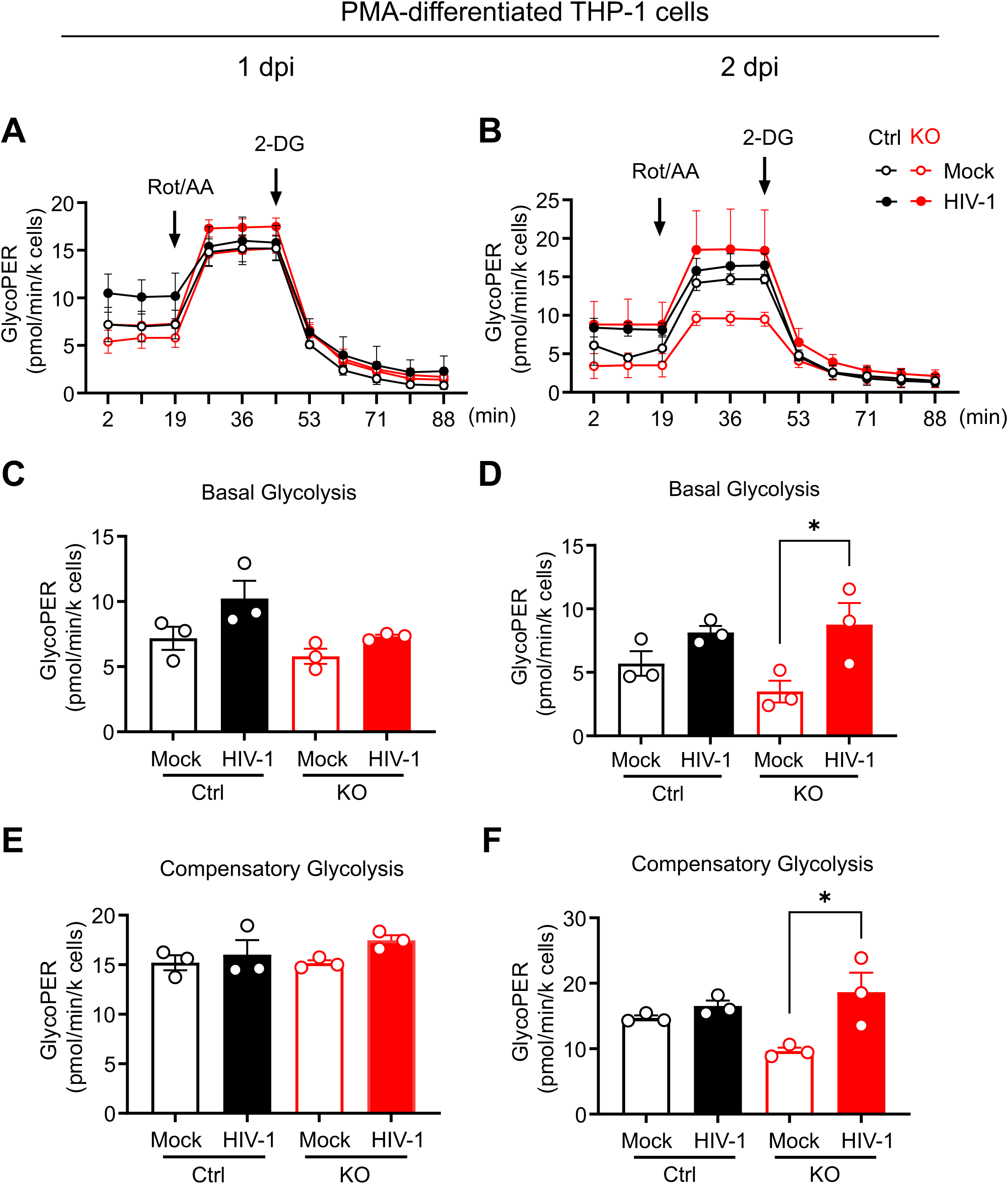
SAMHD1 does not affect glycolysis in differentiated THP-1 cells upon HIV-1 infection. PMA-differentiated THP-1 Ctrl and SAMHD1 KO cells were infected with HIV-1-Luc/VSV-G (MOI = 2) or mock infected. Cells were harvested at 1 dpi (A, C, E) or at 2 dpi (B, D, F) for analyses. (A, B) GlycoPER was measured by a Seahorse XFe24 analyzer at indicated time points and used to evaluate the basal and compensatory glycolysis levels. (C-F) The average values of basal glycolysis (C, D) and compensatory glycolysis (E, F) levels of THP-1 ctrl cells and SAMHD1 KO cells with HIV-1 infection or mock infected. Data are presented as means ± SEM from three independent experiments. (C-F) The one-way ANOVA was used for statistical significance. \**P* < 0.05.

### SAMHD1 does not affect mitochondrial respiration in HIV-1-infected THP-1 cells

Our previous study showed that SAMHD1 enhances HIV-1-induced mitochondrial membrane damage in THP-1 cells and lead to apoptosis (35). Mitochondria are the respiration center of eukaryotic cells (34). To investigate the functional role of SAMHD1 in mitochondrial respiration in monocytic cells during HIV-1 infection, THP-1 Ctrl cells and SAMHD1 KO cells or PMA-differentiated counterparts were infected with HIV-1-Luc/VSV-G. The oxygen consumption rate (OCR) was measured with a Seahorse XFe24 analyzer at 1 and 2 dpi and used to evaluate the mitochondrial respiration levels. The average values of OCR represent basal respiration, ATP production, and maximal respiration. The results of OCR indicated that basal respiration, ATP production, and maximal respiration were not affected by HIV-1 infection or SAMHD1 expression in THP-1 cells at 1 dpi and 2 dpi (Fig. S2). Similar results were observed in PMA-differentiated THP-1 cells as the basal respiration, ATP production, and maximal respiration were not affected by HIV-1 infection or SAMHD1 expression at 1 and 2 dpi (Fig S3). Together, these results suggest that endogenous SAMHD1 does not functionally affect mitochondrial respiration in dividing or differentiated THP-1 cells during HIV-1 infection.

### SAMHD1 enhances HIV-1-induced HK2 expression in THP-1 cells, but not in differentiated cells

To investigate the mechanisms by which SAMHD1 enhances HIV-1-induced basal glycolysis, THP-1 Ctrl cells and SAMHD1 KO cells or PMA-differentiated counterparts were infected with HIV-1 and treated with or without NVP at 1 and 2 dpi. Using Western blotting (WB) and quantification, we measured the expression levels of key glycolytic proteins, including HK1/2, 6-phosphofructo-2-kinase/fructose-2,6-biphosphatase 3 (PFKFB3), phosphofructokinase (PFK), glyceraldehyde-3-phosphate dehydrogenase (GAPDH), pyruvate kinase M1 isoform and M2 isoform (PKM1/2), lactate dehydrogenase (LDH), and pyruvate dehydrogenase (PDH), as well as HIV-1 Gag and capsid (CA) proteins (Figs. S4 and S5) (27). Among the detected cellular proteins, HK1/2 convert glucose into glucose-6-phosphate and drive downstream metabolic cascades (Fig. S5A). We observed that HK1 protein level was lower in Ctrl cells compared with SAMHD1 KO cells, while HK2 protein was generally higher in Ctrl cells compared with SAMHD1 KO cells (Fig. 4A-C, and Fig. S4A). Importantly, HIV-1 infection significantly promoted HK2 expression in Ctrl cells, but not in SAMHD1 KO cells (Fig 4A, 4C and Fig S4A). NVP treatment did not significantly affect HIV-1-induced HK2 expression in THP-1 Ctrl cells. These results suggest that HIV-1 induces HK2 protein expression in THP-1 cells through SAMHD1 independently of productive HIV-1 replication (Fig 4A, 4C and Fig S4A). Moreover, the expression levels of PFKFB3, PFK, GAPDH, PKM1/2, LDH, and PDH were not affected by SAMHD1 expression, HIV-1 infection, or NVP treatment (Fig. S5B). These results suggest that SAMHD1-enhanced basal glycolysis is associated with increased HK2 expression. In differentiated cells, we also observed lower HK1 protein expression in Ctrl cells compared with SAMHD1 KO cells, consistent with the findings in dividing THP-1 cells. However, the expression of HK2 and other key glycolytic proteins was not significantly affected by HIV-1 infection, SAMHD1 expression, or NVP treatment (Fig. 4D-F, and Fig. S5C). Together, these data suggest that endogenous SAMHD1 enhances HIV-1-induced HK2 expression in THP-1 cells, but not in differentiated cells.

**Fig. 4.**
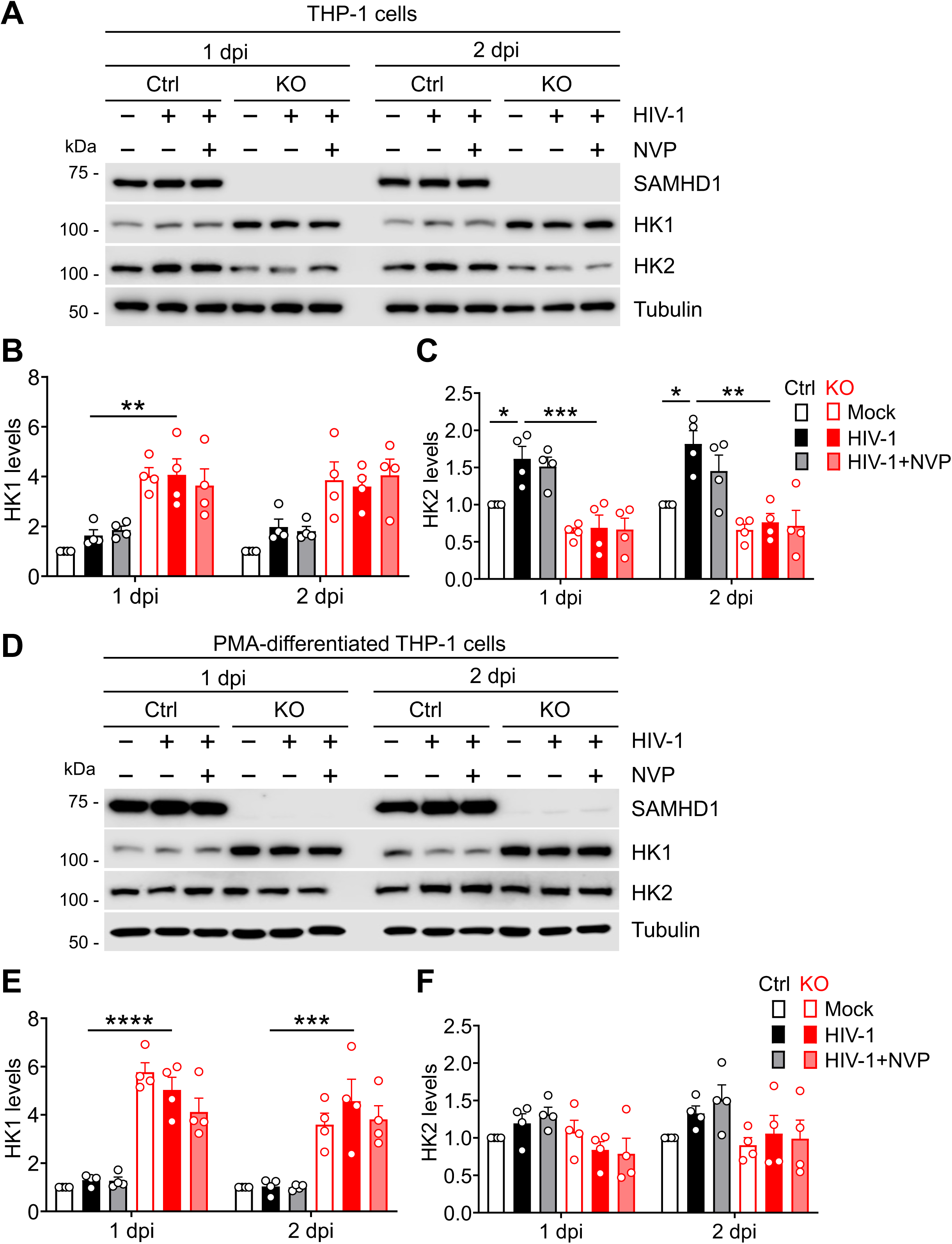
SAMHD1 enhances HIV-1-induced HK2 expression in THP-1 cells, not in differentiated cells. (A-C) THP-1 Ctrl and SAMHD1 KO cells, (D-F) PMA-differentiated THP-1 Ctrl and SAMHD1 KO cells were infected with HIV-1-Luc/VSV-G (MOI= 2) or mock infected. NVP was used to block HIV-1 reverse transcription. The expression levels of SAMHD1, HK1, HK2, and tubulin at 1 and 2 dpi were measured by Western blotting. The relative levels of HK1 (B, E) or HK2 (C, F) protein were quantified by densitometry and normalized to the loading control of tubulin. (A, D) Western blots of one representative are shown. The original Western blots of additional three independent experiments are shown in Fig. S4A. (B, C, E, F) Data are presented as means ± SEM from four independent experiments. HK1 or HK2 protein levels of THP-1 Ctrl cells without HIV-1 infection were set to 1. One-way ANOVA was used for statistical significance. \**P* < 0.05, \*\**P* < 0.01, \*\*\**P* < 0.001, \*\*\*\**P* < 0.0001.

### SAMHD1 enhances HIV-1-induced accumulation of cytosolic HK2 in THP-1 cells, but not in differentiated cells

HK1 and HK2 bind to voltage-dependent anion channels (VDACs) and localize to mitochondria (40). The dissociation of HK1 and HK2 from the mitochondria promotes aerobic glycolysis (41). HK1 and HK2 dissociation from the mitochondria also triggers cytochrome C release and induces apoptosis (40, 42, 43). Our previous study showed that SAMHD1 enhances HIV-1-induced cytochrome C release and apoptosis in THP-1 cells (35). Previous studies showed that SAMHD1 localizes to the mitochondria, where it interacts with both mitochondrial-antiviral signaling protein (MAVS) and VDAC1 (20, 44). Therefore, it is possible that SAMHD1 regulates the mitochondrial recruitment and localization of HK1 and HK2. To explore this possibility, THP-1 Ctrl cells and SAMHD1 KO cells were infected with HIV-1 and treated with or without NVP at 2 dpi and the proteins in cytosolic and mitochondrial fractions were measured by WB (Fig. 5A, 5D, and Fig. S6). Tubulin and VDAC were used as cytosolic and mitochondrial markers, respectively (35). Intriguingly, higher cytosolic HK2 level was detected in HIV-1-infected Ctrl cells with or without NVP treatment compared with in infected SAMHD1 KO cells (Fig. 5A, 5B and Fig. S6A), suggesting that SAMHD1 enhances HIV-1-induced accumulation of cytosolic HK2 in THP-1 cells, which was independent on HIV-1 replication. Conversely, mitochondrial HK2 level was not significantly affected by HIV-1 infection, SAMHD1 expression, or NVP treatment (Fig. 5A, 5C and Fig. S6A). The dissociation of HK1 from mitochondria was not observed in THP-1 cells (Fig. 5A and Fig. S6A).

**Fig. 5.**
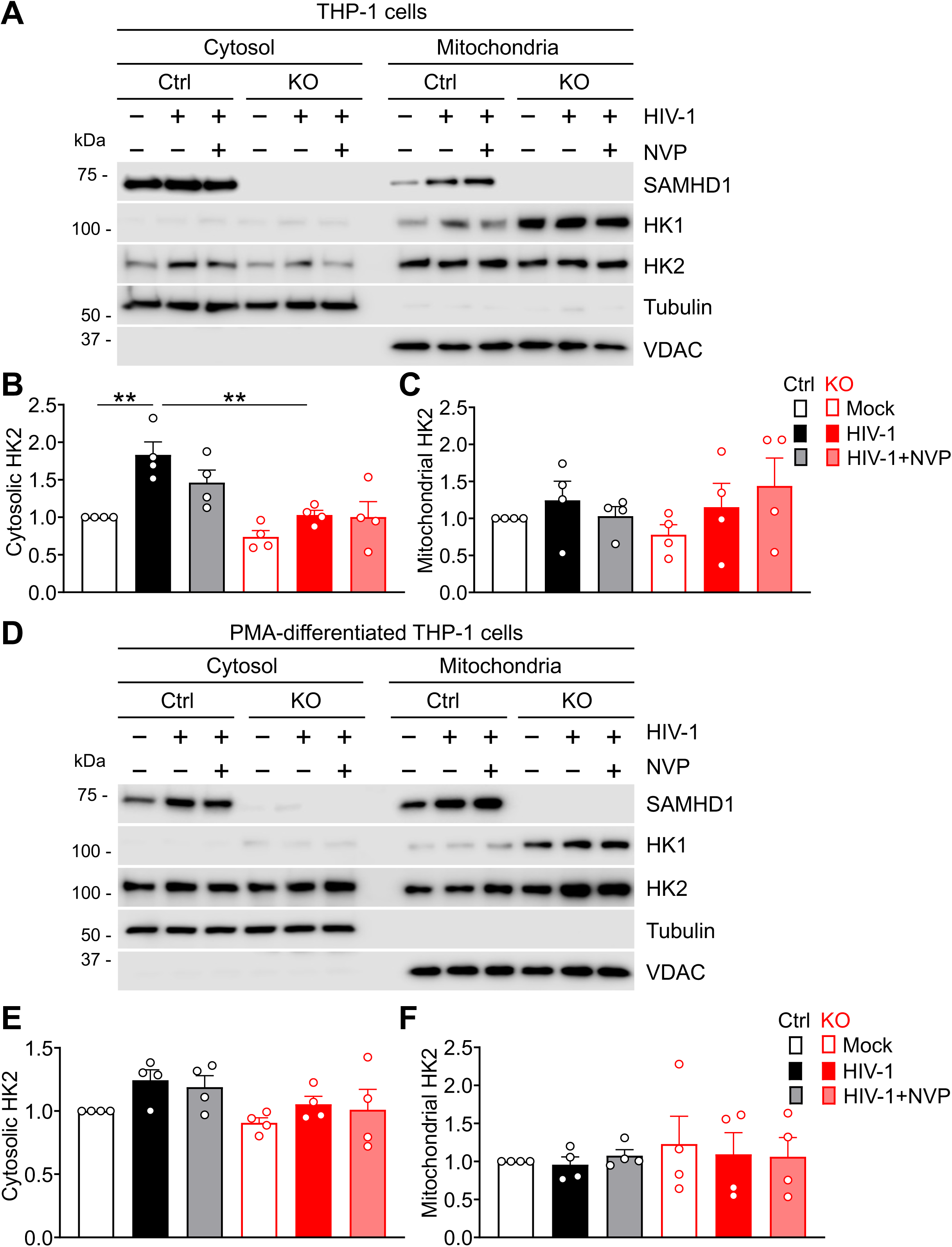
SAMHD1 enhances the accumulation of cytosolic HK2 induced by HIV-1 infection in THP-1 cells, not in differentiated cells. (A-C) THP-1 Ctrl and SAMHD1 KO cells, (D-F) PMA-differentiated THP-1 Ctrl and SAMHD1 KO cells were infected with HIV-1-Luc/VSV-G (MOI= 2) or mock infected. NVP was used to block HIV-1 reverse transcription. The expression levels of SAMHD1, HK1, HK2, VDAC, and tubulin in the cytosolic and mitochondrial fractions at 2 dpi were measured by Western blot. (B, C, E, F) The relative levels of cytosolic HK2 (B, E) and mitochondrial HK2 (C, F) protein were quantified by densitometry and normalized to tubulin and VDAC, respectively. (A, D) Western blots of one representative are shown. The original Western blots of additional three independent experiments are shown in Fig. S4B. (B, C, E, F) Data are presented as means ± SEM from four independent experiments. Cytosolic HK2 or mitochondrial HK2 levels of THP-1 Ctrl cells without HIV-1 infection were set to 1. One-way ANOVA was used for statistical significance. \*\**P* < 0.01.

Our previous study demonstrated that SAMHD1 does not enhance HIV-1-induced apoptosis in PMA-differentiated THP-1 or monocytic U937 cells (35). To explore the effects of SAMHD1 on HK1 and HK2 disassociation from the mitochondria in differentiated THP-1 cells during HIV-1 infection, differentiated Ctrl cells and SAMHD1 KO cells were infected with HIV-1 and treated with or without NVP. At 2 dpi, the levels of cytosolic and mitochondrial HK2 were not affected by HIV-1 infection, SAMHD1 expression, or NVP treatment (Fig. 5D-F, and Fig. S6B). The accumulation of cytosolic HK1 from mitochondria was not observed in differentiated cells (Fig. 5D and Fig. S6B). Together, these data suggest that endogenous SAMHD1 enhances the accumulation of cytosolic HK2 from the mitochondria induced by HIV-1 infection of THP-1 cells, but not in differentiated cells.

### SAMHD1 enhances HIV-1-induced cellular ROS in THP-1 cells, but not in differentiated cells

Cytosolic release of HK2 from the mitochondria increases cellular ROS production and thereby triggers apoptosis or inflammatory responses to infection or inflammation stimuli (42, 45). HIV-1 infection induces ROS production in CD4⁺ T cells and monocytic cells (46–48). To examine the effects of SAMHD1 on cellular ROS production in THP-1 cells upon HIV-1 infection, Ctrl cells and SAMHD1 KO cells or differentiated counterparts were infected with HIV-1 and treated with or without NVP, and cellular ROS levels at 1-4 dpi were measured by H2DCFDA staining and flow cytometry. We observed that cellular ROS levels were higher in HIV-1-infected Ctrl cells compared with infected SAMHD1 KO cells at 1 and 2 dpi (Fig. 6A and Fig. S7A), suggesting that SAMHD1 enhances HIV-1-induced ROS production in THP-1 cells. NVP treatment did not reduce ROS production in HIV-1-infected THP-1 cells, indicating that SAMHD1-enhanced and HIV-1-induced ROS is independent of productive HIV-1 replication. Conversely, cellular ROS levels were not affected by HIV-1 infection, SAMHD1 expression, or NVP treatment in differentiated cells at 1 and 2 dpi (Fig. 6B and Fig. S7B). Thus, endogenous SAMHD1 enhances HIV-1-induced ROS production in THP-1 cells, but not in differentiated cells.

**Fig. 6.**
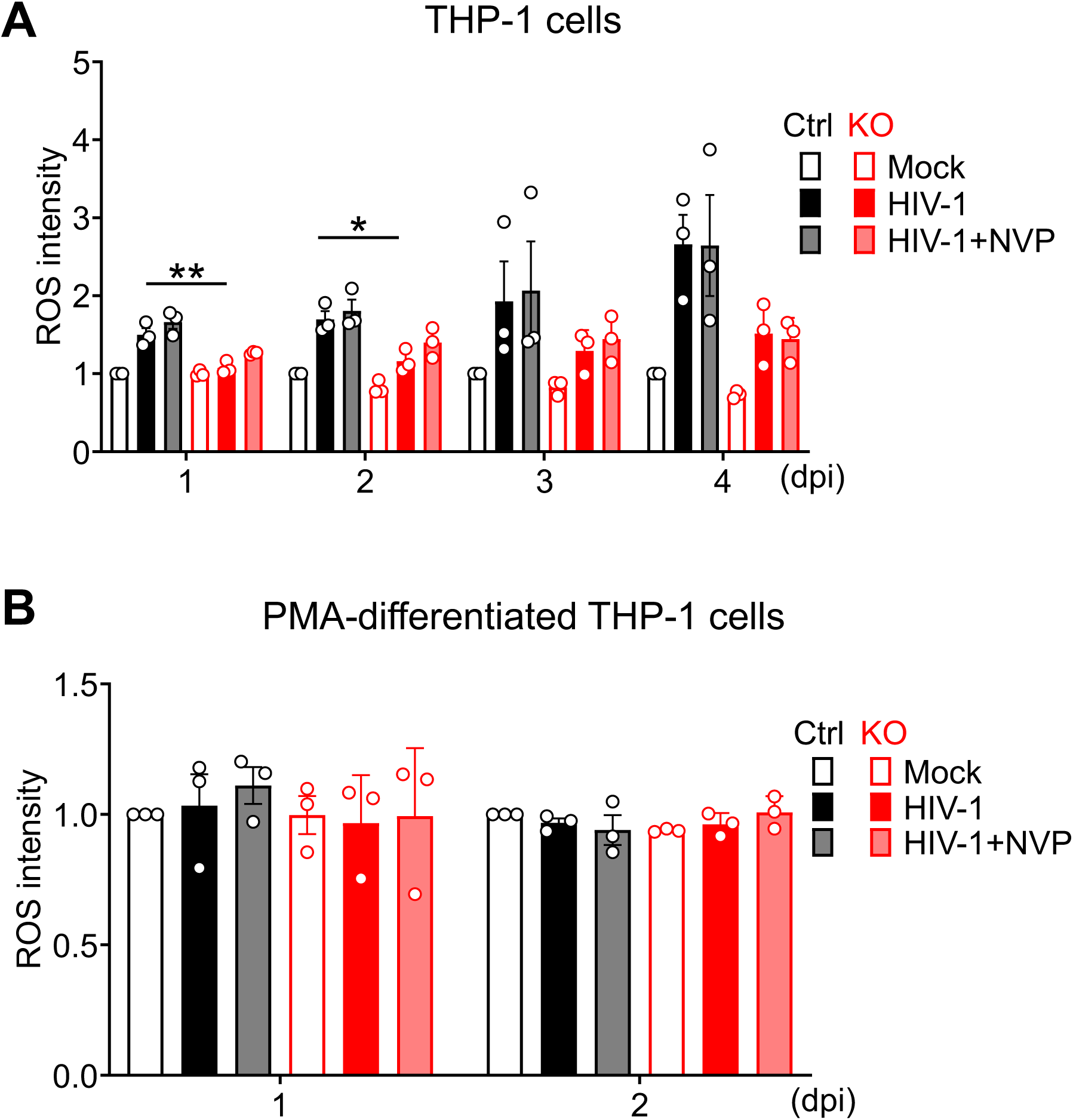
SAMHD1 enhances HIV-1-induced ROS production in THP-1 cells, not in differentiated cells. (A) THP-1 Ctrl and SAMHD1 KO cells, (B) PMA-differentiated THP-1 Ctrl and SAMHD1 KO cells were infected with HIV-1-Luc/VSV-G (MOI = 2) or mock infected. NVP was used to block HIV-1 reverse transcription. THP-1 cells were harvested at 1-4 dpi and PMA-differentiated THP-1 cells were harvested at 1-2 dpi for analyses. ROS was measured by H2DCFDA staining and flow cytometry. The mock levels in Ctrl groups were set as 1. Original flow cytometry results of ROS are shown in Fig. S7. Data are presented as means ± SEM from three independent experiments. The one-way ANOVA was used for statistical significance compared with THP-1 Ctrl cells with HIV-1 infection. \**P* < 0.05, \*\**P* < 0.01.

## DISCUSSION

Glucose metabolism is essential for a wide range of cellular processes, including cellular growth, biosynthesis, immune responses, and cell differentiation (31, 49–51). Glycolysis not only generates ATP but also provides metabolic intermediates required to support these cellular processes (51). In this study, we found that SAMHD1 enhances HIV-1-induced glycolysis in THP-1 cells by increasing HK2 expression and promoting its cytosolic accumulation (Fig. 7). A recent study showed that SAMHD1 knockdown reduces glycolysis and increases mitochondrial stress in monocyte-derived dendritic cells from healthy blood donors (^25^). Other studies showed that SAMHD1 deficiency suppresses glycolytic metabolism while promoting mitochondrial OXPHOS in bone marrow-derived macrophages from mice (^24^). These studies suggest that SAMHD1 is closely linked to cellular metabolism and plays an important role in regulating glucose metabolism in human and mouse myeloid cells. Interestingly, HBV infection induces glucose uptake in target cells and activates the hexosamine biosynthesis pathway, leading to increased O-GlcNAcylation of SAMHD1, which stabilizes SAMHD1 by inhibiting K48-linked ubiquitination and enhances its antiviral activity against HBV (^52^). Therefore, SAMHD1 and glucose metabolism form a bidirectional regulatory loop, in which SAMHD1 influences cellular metabolic state, while glucose metabolism regulates SAMHD1 stability and antiviral function.

**Fig. 7.**
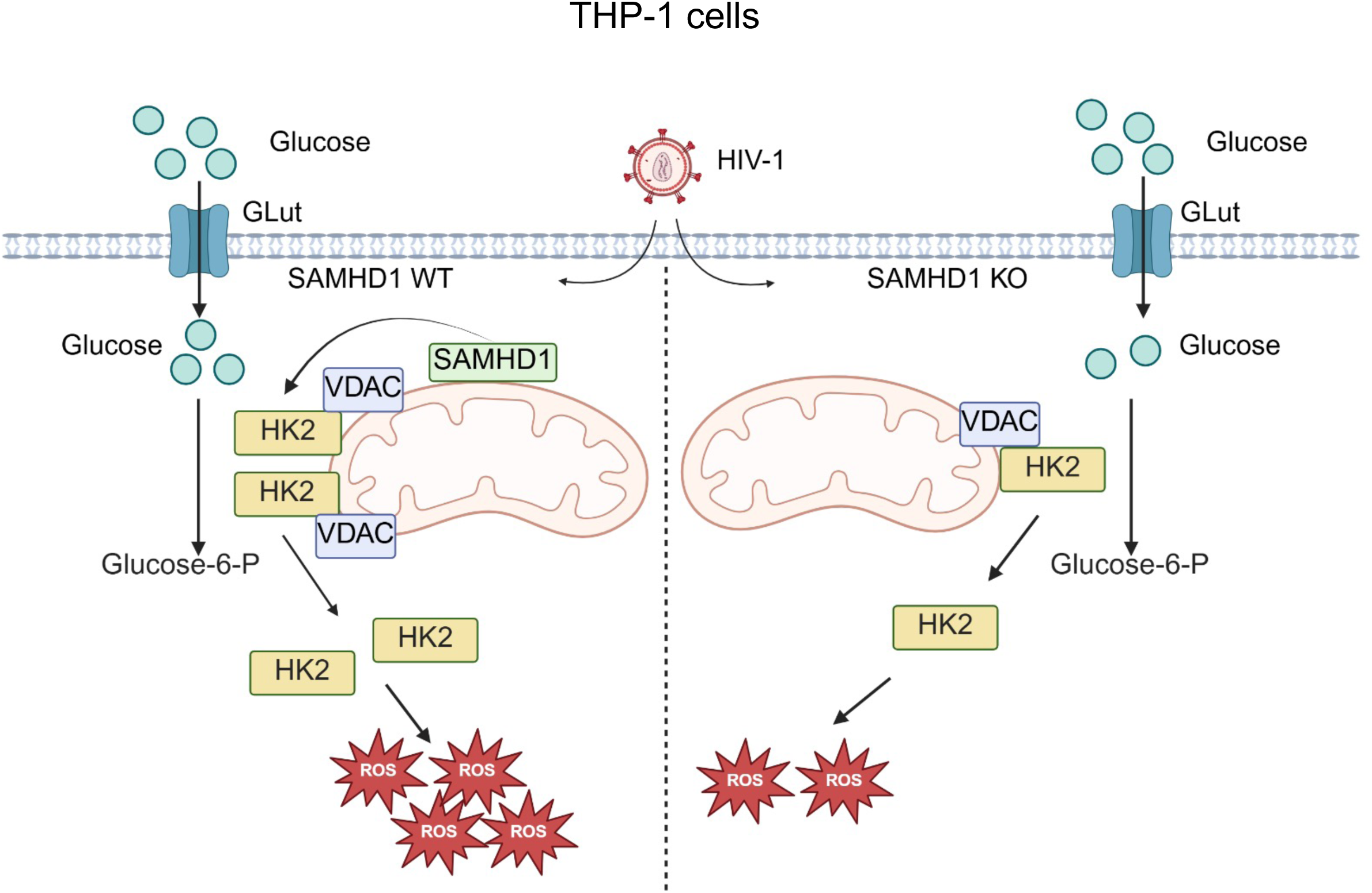
SAMHD1 enhances HIV-1-induced glycolysis and ROS production in THP-1 cells through HK2 upregulation. Endogenous SAMHD1 enhances HIV-1-induced basal glycolysis in dividing THP-1 cells by increasing HK2 expression and promoting cytosolic accumulation of HK2. Moreover, SAMHD1 enhances HIV-1-induced glucose uptake and ROS production in dividing THP-1 cells. These phenotypes were not observed in THP-1-differentiated macrophage-like cells. Our findings suggest that SAMHD1 enhances HIV-1-induced glucose metabolism through a cell state-specific mechanism. This figure was created with BioRender.

SAMHD1 inhibits HIV-1 replication during the early stage of infection but does not suppress HIV-1 infection in THP-1 cells at the late stage of infection. In contrast, SAMHD1 inhibits HIV-1 infection in THP-1-differentiated macrophage-like cells (Fig. 1). Similar data were observed in our previous studies (^22, 35^). This differential effect can be explained by two co-existing scenarios. First, significantly reduced dNTP pools in non-dividing, SAMHD1-high macrophage-like cells are insufficient to support HIV-1 reverse transcription, whereas higher dNTP levels in proliferating, SAMHD1-low monocytic cells reach the threshold required for efficient reverse transcription. Furthermore, phosphorylation of SAMHD1 at T592 negatively regulates its antiviral activity in proliferating monocytic cells, but this regulatory effect is alleviated upon differentiation into non-dividing macrophage-like cells (53). Second, SAMHD1 enhances HIV-1-induced glucose metabolism, which may facilitate viral infection in dividing monocytic cells but not in fully differentiated macrophages.

HIV-1 susceptibility has been shown to positively correlate with glucose metabolic activity in naïve and differentiated memory CD4^+^ T cell subsets (32). HIV-1 infection in primary CD4^+^ T cells and CD4^+^ T cell lines depends on Glut1 upregulation and enhanced glucose uptake (54, 55). In contrast, chronically HIV-1-infected macrophage-like U1 cells, a subclone of U937 cells, exhibit reduced glucose uptake and decreased glycolytic intermediates compared with parental U937 cells (36). Furthermore, HIV-1 infection does not significantly alter glycolytic or TCA cycle intermediates in human primary monocyte-derived macrophages (56). These findings are consistent with our data showing that glucose uptake and glycolysis are induced by HIV-1 infection in dividing THP-1 cells, whereas these metabolic parameters are not significantly affected in differentiated macrophage-like cells. Therefore, HIV-1-induced metabolic reprogramming is highly dependent on cell types. In addition, glycolysis is not only involved in acute HIV-1 infection but also in the regulation of viral latency (57, 58). Previous studies reported that HIV-1 latency is associated with reduced glycolytic activity and increased sensitivity to oxidative stress in CD4^+^ T cells, suggesting that glucose metabolism may contribute to both the maintenance and reactivation of viral latency (57, 58).

A previous study showed that HIV-1 infection in primary human CD4^+^ T cells induces HK1, but not HK2 expression (^33^). The HK1 upregulation is independent of individual HIV-1 accessory proteins, including Vif, Vpr, Vpu, and Nef, suggesting that HK1 induction is not mediated by a single accessory protein but rather reflects broader aspects of HIV-1 infection in primary CD4^+^ T cells (33). In addition, the HIV-1 envelope protein gp120 has been shown to upregulate glycolytic enzymes, including HK1 and enolase 2 in glioma cells (^59^). In our study, treatment with the reverse transcriptase inhibitor NVP completely blocked HIV-1 infection but did not affect HIV-1-induced glucose uptake, HK2 expression, or ROS production. One possible explanation is that single-cycle HIV-1 virions contain several viral proteins, including CA, and Vpr. HIV-1 Vpr can reprogram macrophage metabolism by activating glycolysis, accompanied by increased expression of HK1 and HK2 in U937-derived macrophage-like cells (^60, 61^). Therefore, although NVP efficiently inhibits HIV-1 reverse transcription and infection, it does not affect virion-associated proteins in incoming virions, such as CA (Fig. S5) or Vpr, which might contribute to the observed metabolic effects despite the blockade of viral infection. Moreover, we found that SAMHD1 promoted HIV-induced accumulation of cytosolic HK2 and increased intracellular ROS production in THP-1 cells but not in differentiated THP-1 cells. These effects can further lead to cytochrome c release and subsequent apoptosis in dividing THP-1 cells, but not in differentiated cells as we previously reported (35). Thus, our findings also provide an mechanistic explanation for our previous results that SAMHD1 enhances HIV-1-induced apoptosis in THP-1 cells through the mitochondrial pathway (35).

In summary, we demonstrated that SAMHD1 promotes glycolytic metabolism in HIV-1-infected monocytic cells through HK2 upregulation. Our findings revealed a previously unrecognized function of SAMHD1 in modulating cellular responses to HIV-1 infection.

## MATERIALS AND METHODS

### Key reagents and resources

Please refer to detailed information summarized in Table 1.

**TABLE 1:** KEY REAGENTS AND RESOURCES.

| REAGENT or RESOURCE | SOURCE | IDENTIFIER |
| --- | --- | --- |
| **Antibodies** |  |  |
| Mouse monoclonal anti-SAMHD1 | Abcam | Cat#ab117908;<br>RRID:AB_10901093 |
| Rabbit monoclonal anti-hexokinase I | Cell Signaling Technologies (CST) | Cat#2024;<br>RRID:AB_2116996 |
| Rabbit Monoclonal anti-hexokinase II | CST | Cat#2867;<br>RRID:AB_2232946 |
| Mouse monoclonal anti-tubulin | DSHB | Cat#12G10;<br>RRID:AB_528498 |
| Rabbit monoclonal anti-VDAC | CST | Cat#4661;<br>RRID:AB_10557420 |
| Rabbit monoclonal anti-PFKFB3 | CST | Cat#13123;<br>RRID:AB_2617178 |
| Rabbit monoclonal anti-PFKP | CST | Cat#8164;<br>RRID:AB_2713957 |
| Rabbit monoclonal anti-GAPDH | CST | Cat#5174;<br>RRID:AB_10622025 |
| Rabbit monoclonal anti-PKM1/2 | CST | Cat#3190;<br>RRID:AB_2163695 |
| Rabbit monoclonal anti-PKM2 | CST | Cat#4053<br>RRID:AB_1904096 |
| Rabbit monoclonal anti-pyruvate dehydrogenase | CST | Cat#3205;<br>RRID:AB_2162926 |
| Rabbit monoclonal anti-LDHA | CST | Cat#3582;<br>RRID:AB_2066887 |
| Mouse monoclonal anti-HIV-1 p24 (capsid) | The NIH AIDS Reagent Program | ARP-4121 |
| Goat anti-mouse IgG (H+L) HRP | Promega | Cat#W4021;<br>RRID:AB_430834 |
| Goat anti-rabbit IgG (H+L) HRP | Promega | Cat#W4011;<br>RRID:AB_430833 |
| **Bacterial and virus strains** |  |  |
| Single-cycle, VSV-G-pseudotyped luciferase reporter HIV-1 (based on pNL4-3E-R+) | The NIH AIDS Reagent Program (22) | Generated from pNL4-3E-R+ |
| **Chemicals, peptides, and recombinant proteins** |  |  |
| DMEM | Gibco | 11965-092 |
| RPMI-1640 medium | ATCC | 30-2001 |
| RPMI-1640 medium (no glucose) | Gibco | 11879-020 |
| Fetal bovine serum (FBS) | R&D Systems | S1150H |
| Penicillin and streptomycin | Gibco | 15140-122 |
| Puromycin | Millipore-Sigma | P8833 |
| Nevirapine (NVP) | The NIH AIDS Reagent Program | 4666 |
| Cell lysis buffer | CST | 9803 |
| Protease inhibitor cocktail | Millipore-Sigma | P8340 |
| Phosphatase inhibitor cocktail | Millipore-Sigma | P0044 |
| jetPRIME® transfection reagent | Polyplus | 101000046 |
| Geneticin (G418) | Gibco | 10131035 |
| Phorbol 12-myristate 13-acetate (PMA) | Millipore-Sigma | P8139 |
| 2-NBDG | MCE | HY-116215 |
| Poly-α-Lysine hydrobromide | Millipore-Sigma | P9155 |
| Seahorse XF RPMI medium, pH 7.4 | Agilent | 103576-100 |
| Rotenone (Rot) | Millipore-Sigma | R8875 |
| Antimycin A (AA) | Millipore-Sigma | A8674 |
| 2-Deoxy-D-glucose (2-DG) | Thermo Fisher Scientific | 111980050 |
| Oligomycin A (Oligo) | Millipore-Sigma | 75351 |
| FCCP (Carbonyl cyanide 4-(trifluoromethoxy) phenylhydrazone) | Millipore-Sigma | C2920 |
| Hoechst 33342 | Thermo Fisher Scientific | 62249 |
| Critical commercial assays |  |  |
| Luciferase Assay System | Promega | E1500 |
| Cell Fractionation Kit - Standard | Abcam | ab109719 |
| Pierce™ BCA Protein Assay Kits | Thermo Fisher Scientific | 23225 |
| Cytiva Amersham ECL Prime Western Blotting Detection Reagent | Cytiva | 45002401 |
| SuperSignal West Femto Maximum Sensitivity Substrate | Thermo Fisher Scientific | 34094 |
| DCFDA / H2DCFDA - Cellular ROS Assay Kit | abcam | ab113851 |
| Universal mycoplasma detection kit | ATCC | 30-101-2K |
| Experimental models: Cell lines |  |  |
| THP-1 control (Ctrl) and THP-1 SAMHD1 knockout (KO) cells | The Wu lab (22) | N/A <sup>a</sup> |
| GHOST/R5/X4 cell line | Vineet KewalRamani(66) | N/A <sup>a</sup> |
| HEK 293T cell line | ATCC | CRL-3216 |
| Recombinant DNA |  |  |
| pNL4-3E-R+ Luciferase | The NIH AIDS Reagent Program(18) | N/A <sup>a</sup> |
| pMD2.G | Addgene | Cat#12259 |
| Software and algorithms |  |  |
| Flowjo V10 | BD Biosciences | FlowJo licensed user |
| GraphPad Prism 10 | GraphPad | Prism – GraphPad licensed user |
| BioRender | BioRender.com | BioRender licensed user |

<sup>a</sup> N/A, not available.

### Cell culture

THP-1 Ctrl and THP-1 SAMHD1 KO cell lines were described previously (22). The cells were cultured in RPMI 1640 (ATCC) with 10% fetal bovine serum (FBS), 100 U/mL penicillin, 100 μg/mL streptomycin, and 1 μg/mL puromycin. HEK293T cells and GHOST/R5/X4 cells were described (18). HEK293T cells were cultured in DMEM with 10% FBS, 100 U/mL penicillin, and 100 μg/mL streptomycin. GHOST/R5/X4 were cultured in DMEM with 10% FBS, 100 U/mL penicillin, 100 μg/mL streptomycin, 1 μg/mL puromycin, 500 μg/mL G418 and 50 μg/mL hygromycin B. All cell lines were cultured at 37°C in a humidified atmosphere containing 5% CO_2_ and were confirmed to be free of mycoplasma contamination using a universal mycoplasma detection kit.

### HIV-1 stocks and viral infection assays

Single-cycle HIV-1-Luc pseudotyped with VSV-G was generated by co-transfecting HEK293T cells with pNL4-3E-R+ Luciferase and pMD2.G. Virus stock was digested with DNase I (40 U/mL) for 1 h at 37°C and subsequently filtered through a 0.45 μm filter. Viral infectivity was determined by limiting dilution on GHOST/X4/R5 cells as previously described (62). THP-1 cells or PMA-differentiated THP-1 cells were infected with HIV-1-Luc/VSV-G at a multiplicity of infection (MOI) of 2 in the presence of 10 μg/mL polybrene. HIV-1 infection was quantified by luciferase assay according to the manufacturer’s instructions. Luciferase activity was normalized to total protein concentration measured using a bicinchoninic acid (BCA) assay kit.

### Treatment of cells with NVP or PMA

THP-1 cell lines were infected with HIV-1-Luc/VSV-G. Nevirapine (NVP, 10 μM) was maintained in the medium throughout the infection and subsequent culture period. For PMA differentiation, THP-1 cells were treated with PMA (30 ng/mL) for 24 h and then cultured in fresh medium for an additional 24 h before further analyses or experiments.

### Measurement of glucose uptake

THP-1 cells or PMA-differentiated THP-1 cells were infected with HIV-1-Luc/VSV-G in the presence or absence of NVP. Cells were incubated with 50 μM 2-NBDG in glucose-free RPMI 1640 medium for 1 h (63, 64). After incubation, cells were washed with PBS, and glucose uptake was measured by flow cytometry.

### Measurement of GlycoPER and OCR

GlycoPER and OCR were measured using a Seahorse XF24 Extracellular Flux Analyzer according to the manufacturer’s instructions. HIV-1-infected or mock-infected THP-1 cells were seeded at a density of 8 × 10 cells per well in XF24 plates pre-coated with 0.1 mg/mL poly-L-lysine hydrobromide. For PMA-differentiated THP-1 cells, 8 × 10 THP-1 cells were seeded in XF24 plates and differentiated with PMA. Following PMA treatment, cells were infected with HIV-1-Luc/VSV-G. Prior to analysis, cells were incubated in Seahorse XF RPMI Medium for 1 h at 37°C in a non-CO_2_ incubator. The XF24 plates were then loaded into the Seahorse analyzer for metabolic measurements. For the glycolytic rate assay, rotenone (0.5 μM) and antimycin A (0.5 μM) were injected through port A, followed by 2-DG (50 mM) through port B and Hoechst 33342 through port C. For the mitochondrial stress test, oligomycin A (1 μM), FCCP (2 μM), rotenone (1 μM) plus antimycin A (1 μM), and Hoechst 33342 were sequentially injected through ports A, B, C, and D, respectively. After the Seahorse assay, XF24 plates were imaged using an Agilent BioTek Cytation imaging system. The number of Hoechst 33342-positive nuclei was quantified and used for normalization of GlycoPER and OCR measurements, which were reported per 1,000 cells.

### Western blotting

Western blot analysis was performed as previously described (19, 35). Briefly, cells were lysed in cell lysis buffer supplemented with protease inhibitor cocktail and phosphatase inhibitor cocktail. Protein concentrations were determined using a BCA assay. Equal amounts of protein were separated by SDS-PAGE and transferred onto nitrocellulose membranes. Membranes were blocked with 5% non-fat milk for 1 h and incubated with the indicated primary antibodies overnight at 4°C, followed by HRP-conjugated secondary antibodies for 1 h at room temperature. Protein bands were detected using enhanced chemiluminescence (ECL) and visualized with an Odyssey Fc Imager. Tubulin was used as the loading control.

### Cytosolic and mitochondrial fractionation

To examine the subcellular distribution of proteins, THP-1 cells were mock infected or infected with HIV-1-Luc/VSV-G in the presence or absence of NVP for 2 dpi. Cytoplasmic and mitochondrial fractions were prepared using a Cell Fractionation Kit following the manufacturer’s protocol. All buffers were supplemented with protease and phosphatase inhibitor cocktails. Proteins from each fraction were subjected to Western blot analysis. The purity of the fractions was verified using tubulin as a cytoplasmic marker and VDAC as a mitochondrial marker (35, 65).

### Measurement of ROS

THP-1 cells or PMA-differentiated THP-1 cells were infected with HIV-1-Luc/VSV-G in the presence or absence of NVP. Intracellular ROS levels were measured using a Cellular ROS Assay Kit according to the manufacturer’s instructions. Briefly, cells were incubated with 20 μM DCFDA in 1× assay buffer for 30 min at 37°C in the dark. After incubation, cells were washed with 1× assay buffer, and ROS levels were analyzed by flow cytometry.

### Statistical analysis

All data are presented as the mean ± SEM. Statistical analyses were performed using GraphPad Prism software. Two-way analysis of variance (ANOVA) was used to evaluate differences between groups. P value less than 0.05 was considered statistically significant.

## ACKNOWLEDGMENTS

We thank the members of the Wu laboratory for their helpful discussions and suggestions. We appreciate the reagents provided by the National Institutes of Health (NIH) AIDS Reagent Program. We thank Dr. Eric Weatherford of the Metabolic Phenotyping Core at the University of Iowa for his assistance and support. We also thank Drs. Eric Taylor and Ling Yang for critically reading the manuscript and their constructive feedback. This work was supported by NIH grant R01AI189220 (to L.W.). L.W. and his lab are also supported by NIH grants R33AI169659 and R21AI181742. The content is solely the responsibility of the authors and does not necessarily represent the official views of the NIH.

## AUTHOR CONTRIBUTIONS

Conceptualization, H.Y. and L.W.; methodology, H.Y.; validation, H.Y.; formal analysis, H.Y.; investigation, H.Y.; resources, L.W.; visualization, H.Y. and L.W.; funding acquisition, L.W.; writing– original draft, H.Y., P.H.C. and L.W.; writing– review & editing, H.Y., P.H.C. and L.W.

## DECLARATION OF INTERESTS

The authors declare no competing interests.

## SUPPLEMENTAL INFORMATION

Supplemental Figures S1–S7 and legends

## SUPPLEMENTAL FIGURE LEGENDS

**Fig. S1.**
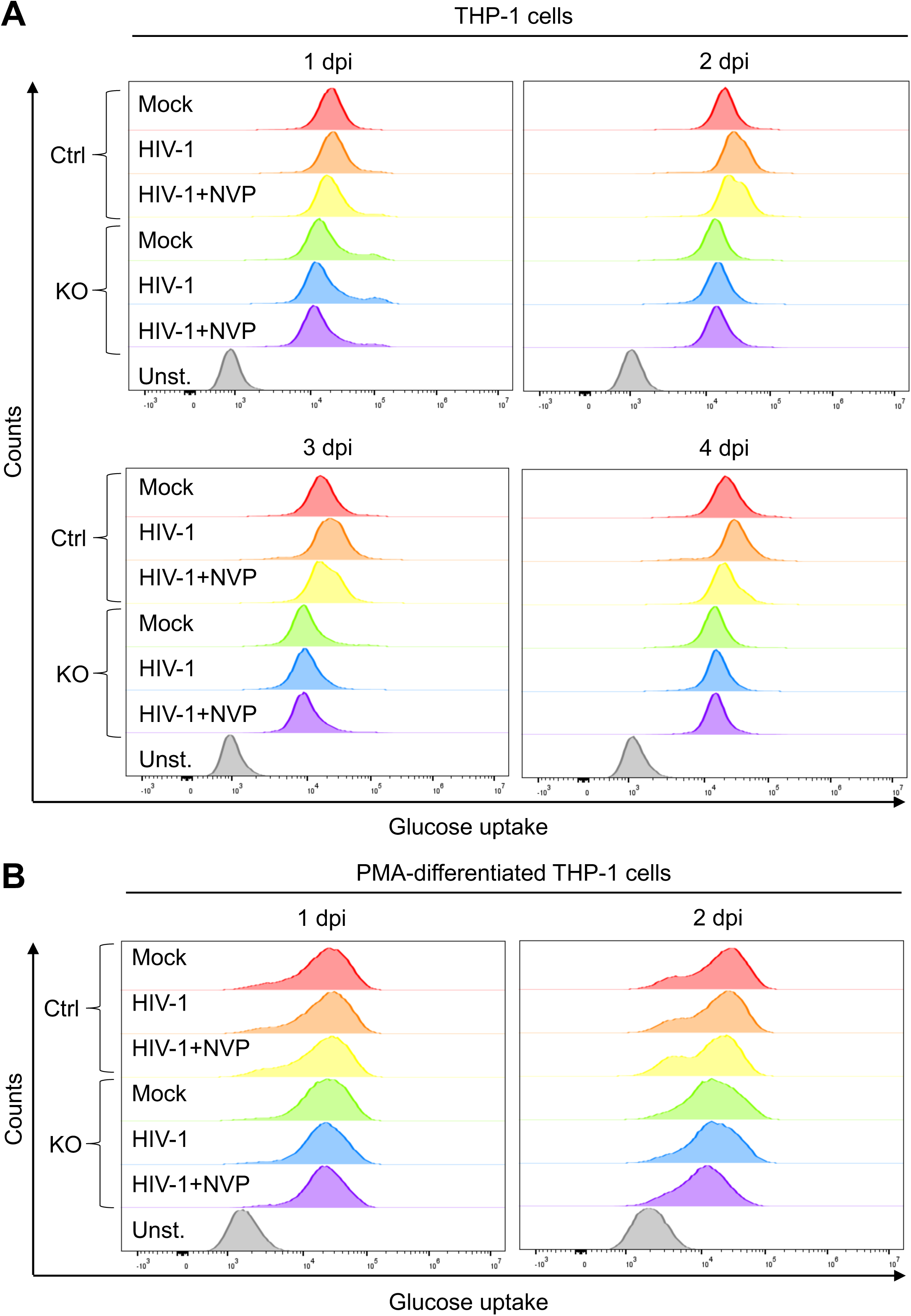
Endogenous SAMHD1 enhances HIV-1-induced glucose uptake in THP-1 cells, not in differentiated cells. (A) THP-1 Ctrl and SAMHD1 KO cells, (B) PMA-differentiated THP-1 Ctrl and SAMHD1 KO cells were infected with HIV-1-Luc/VSV-G (MOI = 2) or mock infected. NVP was used to block HIV-1 reverse transcription. THP-1 cells were harvested at 1-4 dpi and differentiated THP-1 cells were harvested at 1-2 dpi for analyses. Glucose uptake was measured by 2-NBDG staining and flow cytometry. Unst., Unstained cells.

**Fig. S2.**
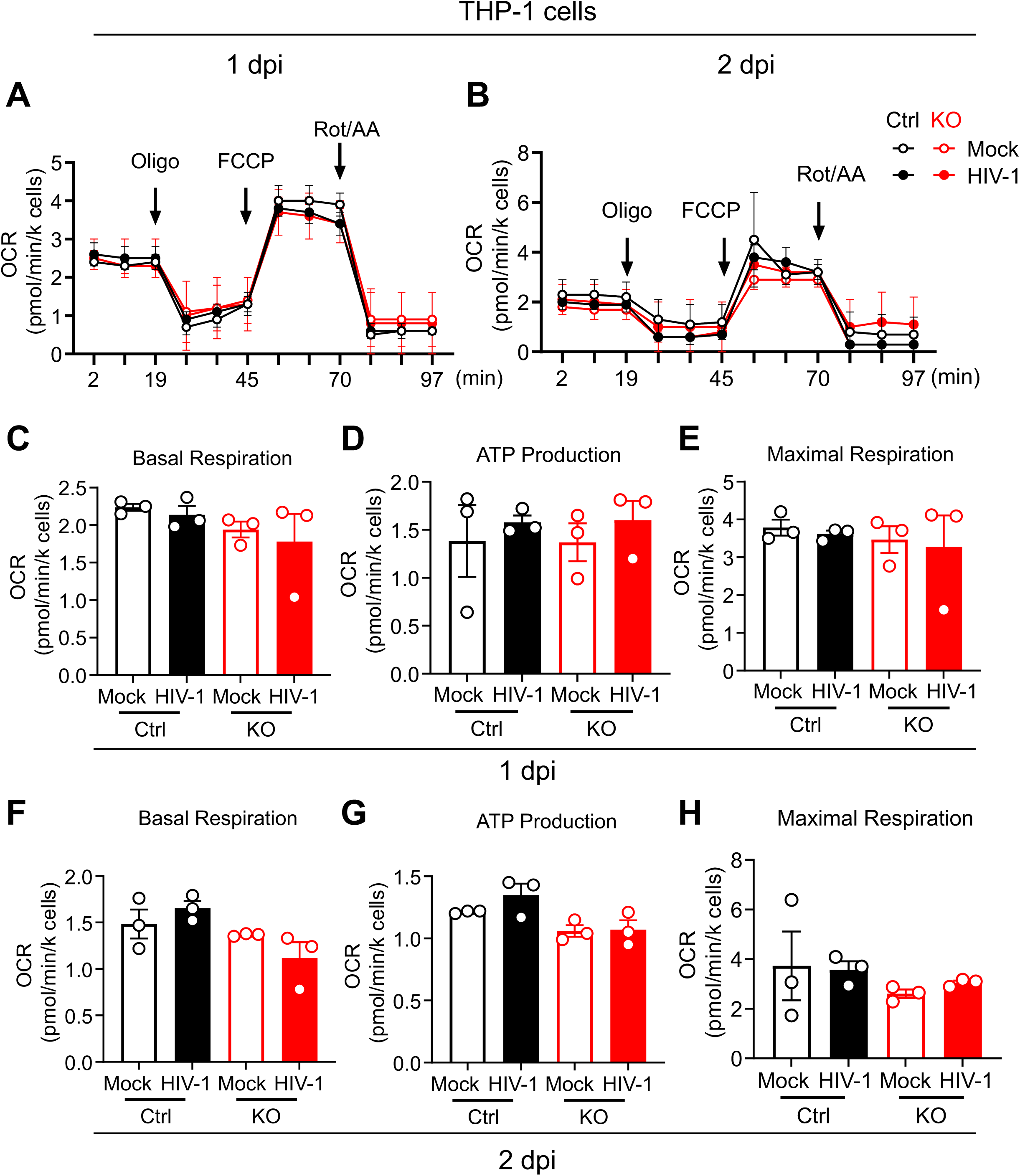
SAMHD1 does not affect mitochondrial respiration in HIV-1-infected THP-1 cells. THP-1 Ctrl and SAMHD1 KO cells were infected with HIV-1-Luc/VSV-G (MOI = 2) or mock infected. Cells were harvested at 1 dpi (A, C, D, E) and 2 dpi (B, F, G, H) for analyses. (A, B) The OCR was measured by a Seahorse XFe24 analyzer at indicated time points and was used to evaluate the mitochondrial respiration level. (C-H) The average values of OCR representing basal respiration (C, F), ATP Production (D, G), and maximal respiration (E, H) of THP-1 Ctrl and SAMHD1 KO cells with HIV-1 infection or mock infected were plotted. Data are presented as means ± SEM from three independent experiments. (C-H) The one-way ANOVA was used for statistical significance (no significance was identified).

**Fig. S3.**
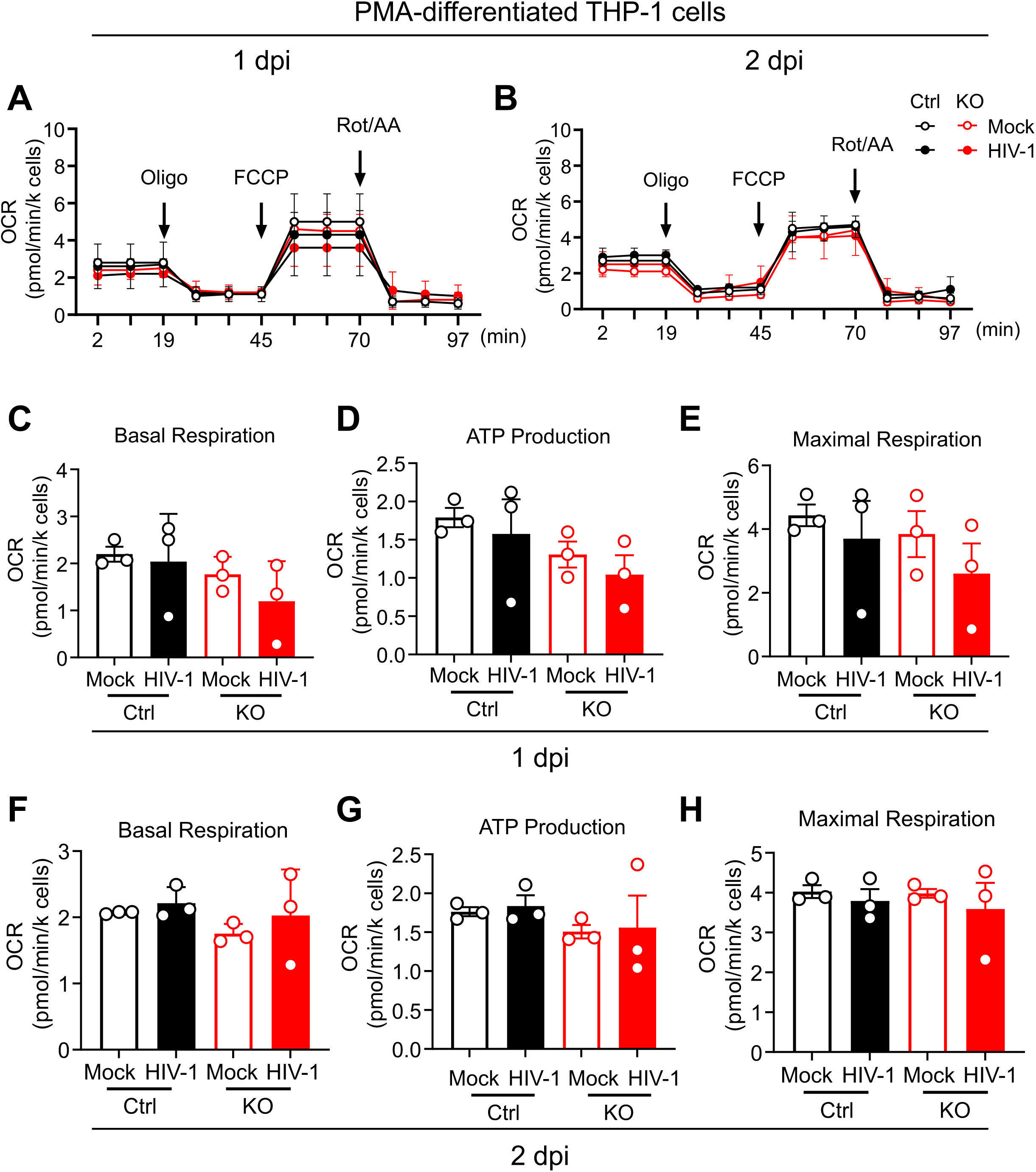
SAMHD1 does not affect mitochondrial respiration in differentiated THP-1 cells infected with HIV-1. PMA-differentiated THP-1 Ctrl and SAMHD1 KO cells were infected with HIV-1-Luc/VSV-G (MOI = 2) or mock infected. Cells were harvested at 1 dpi (A, C, D, E) or 2 dpi (B, F, G, H) for analyses. (A, B) OCR was measured at indicated time points by a Seahorse XFe24 analyzer and used to evaluate the mitochondrial respiration level. (C-H) The average values of OCR representing basal respiration (C, F), ATP Production (D, G) and maximal respiration (E, H) of differentiated THP-1 Ctrl and SAMHD1 KO cells with HIV-1 infection or mock infected were plotted. Data are presented as means ± SEM from three independent experiments. (C-H) The one-way ANOVA was used for statistical significance (no significance was identified).

**Fig. S4.**
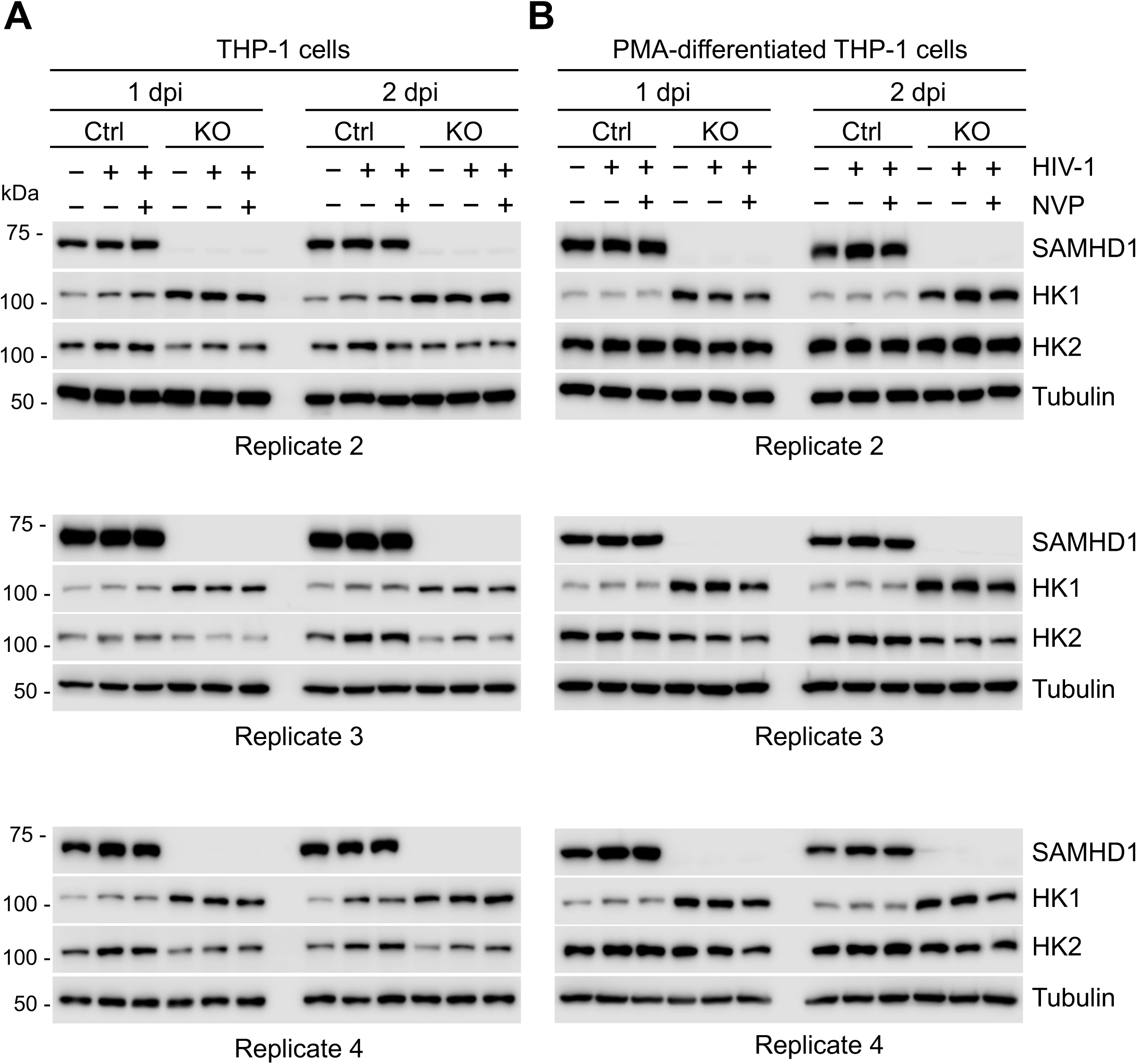
SAMHD1 enhances HIV-1-induced HK2 expression in THP-1 cells, not in differentiated cells. (A) THP-1 Ctrl and SAMHD1 KO cells, (B) PMA-differentiated THP-1 Ctrl and SAMHD1 KO cells were infected with HIV-1-Luc/VSV-G (MOI = 2) or mock infected. NVP was used to block HIV-1 reverse transcription. The expression levels of SAMHD1, HK1, HK2, and tubulin at 1 and 2 dpi were measured by Western blotting. Results of three independent experiments (replicates 2-4) are shown.

**Fig. S5.**
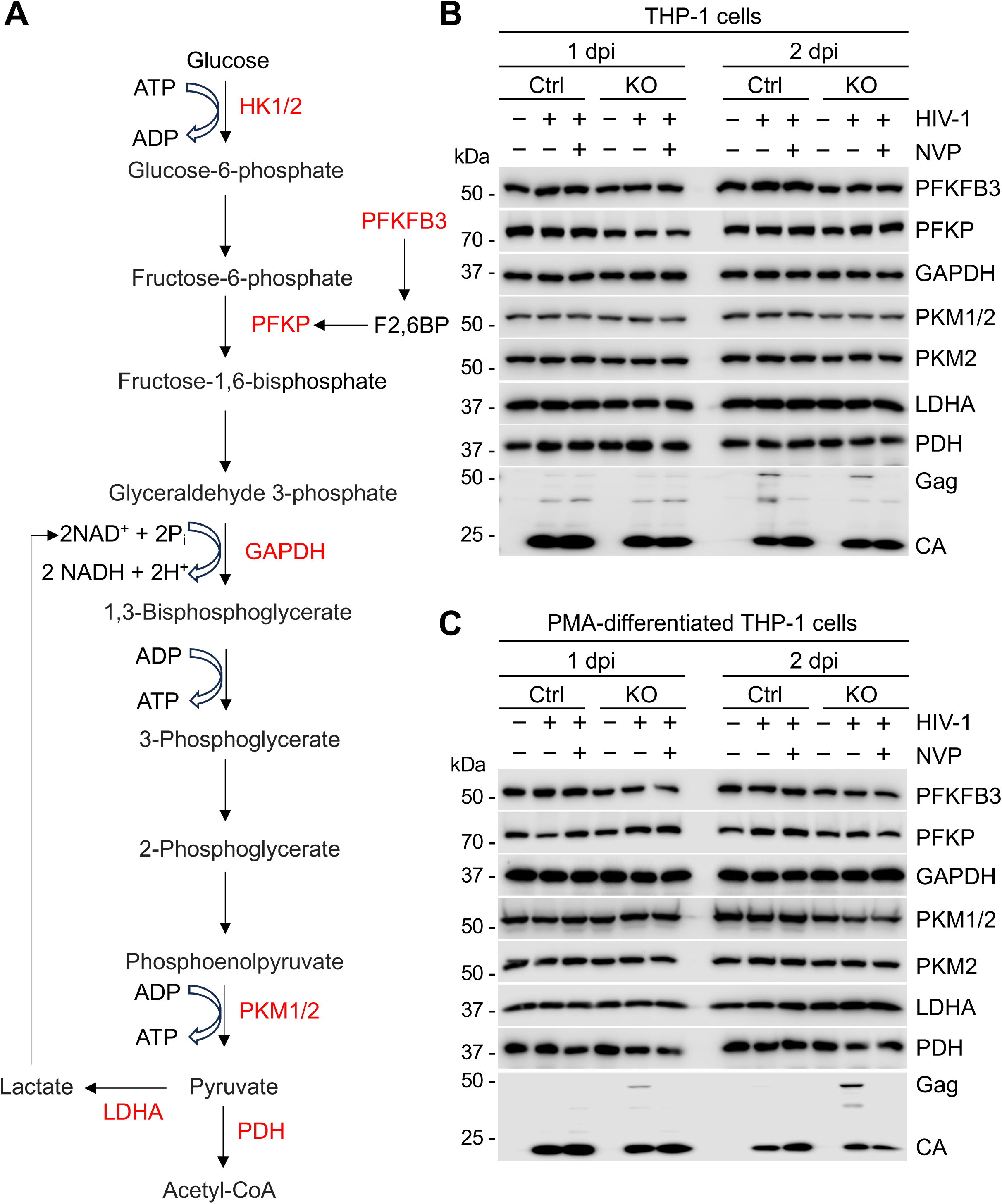
SAMHD1 does not affect the expression of PFKFB3, PFKP, GADPH, PKM1/2, LDHA and PDH in THP-1 cells and differentiated cells upon HIV-1 infection. (A) Schematic overview of the glycolytic pathway with key proteins. The proteins indicated with red fonts were detected by Western blotting in (B and C). (B) THP-1 Ctrl and SAMHD1 KO cells, (C) PMA-differentiated THP-1 Ctrl and SAMHD1 KO cells were infected with HIV-1-Luc/VSV-G (MOI= 2) or mock infected. NVP was used to block HIV-1 reverse transcription. PFKFB3, PFKP, GADPH, PKM1/2, LDHA, PDH, as well as HIV-1 Gag and CA at 1 and 2 dpi were detected by Western blotting.

**Fig. S6.**
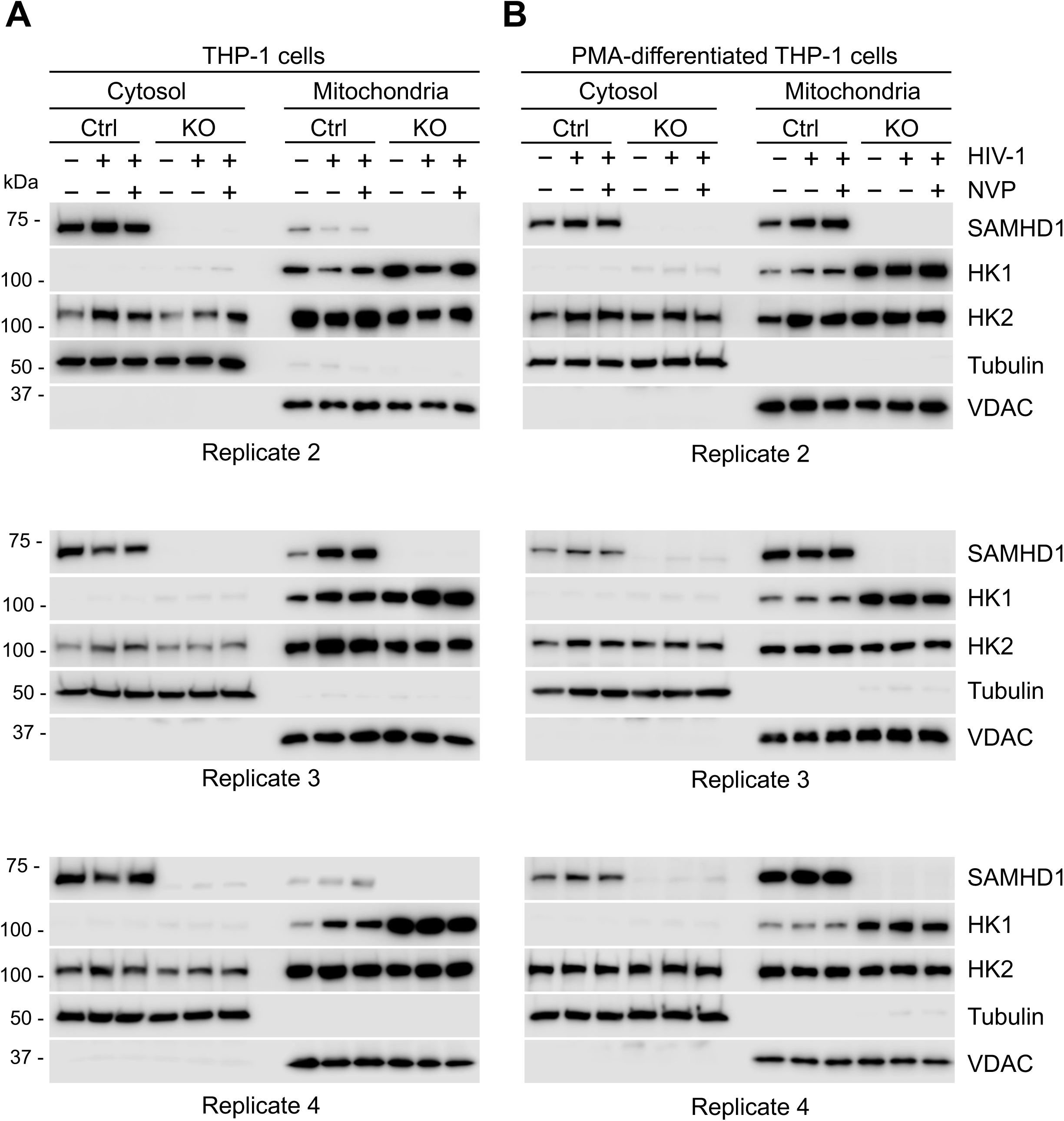
SAMHD1 enhances the accumulation of cytosolic HK2 induced by HIV-1 infection in THP-1 cells, but not in differentiated cells. (A) THP-1 Ctrl and SAMHD1 KO cells, (B) PMA-differentiated THP-1 Ctrl and SAMHD1 KO cells were infected with HIV-1-Luc/VSV-G (MOI= 2) or mock infected. NVP was used to block HIV-1 reverse transcription. SAMHD1, HK1, HK2, VDAC, and tubulin in the cytosolic and mitochondrial fractions at 2 dpi were detected by Western blotting. Results of three independent experiments (replicates 2-4) are shown.

**Fig. S7.**
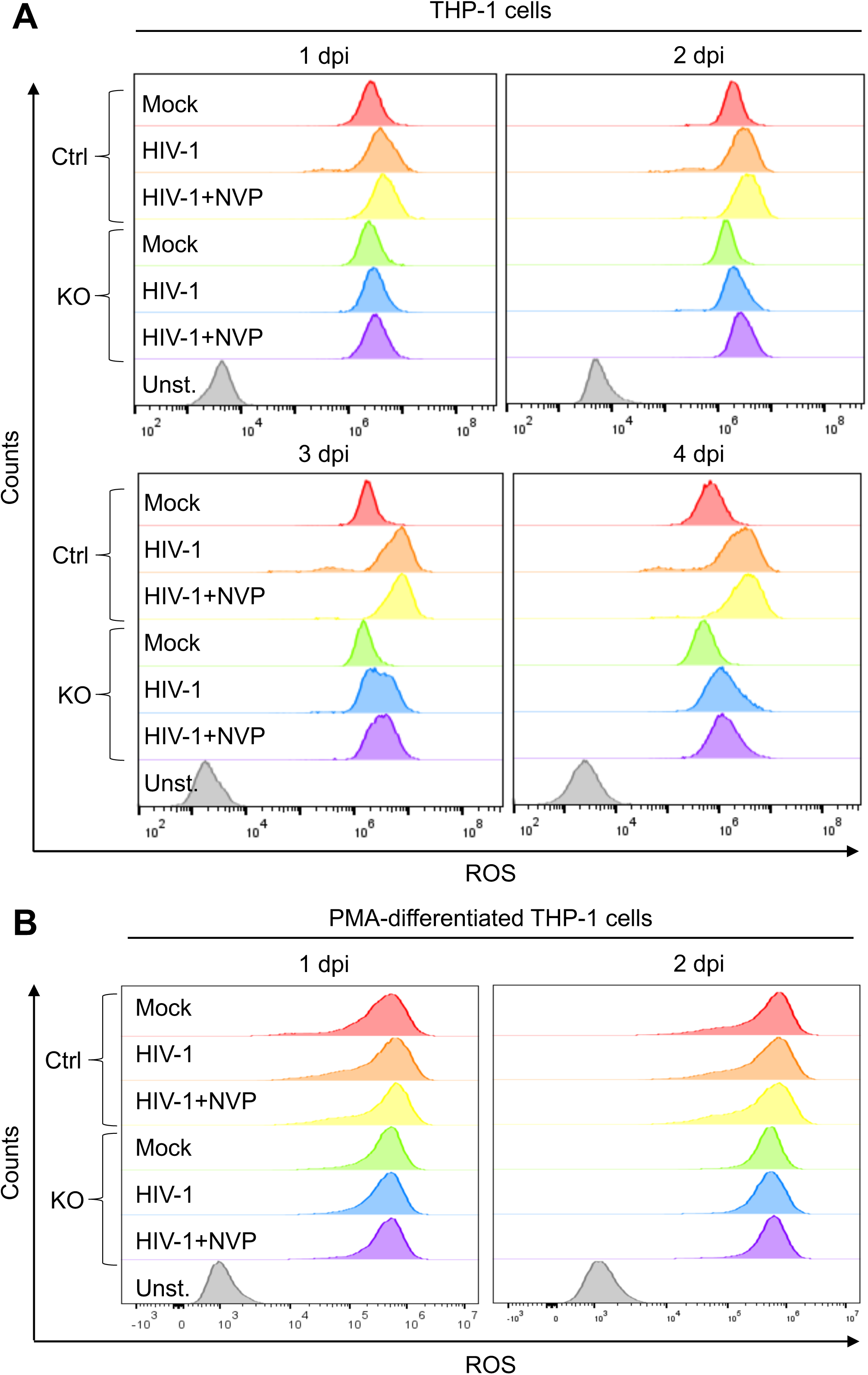
SAMHD1 enhances HIV-1-induced ROS production in THP-1 cells, not in differentiated cells. (A) THP-1 Ctrl and SAMHD1 KO cells, (B) PMA-differentiated THP-1 Ctrl and SAMHD1 KO cells were infected with HIV-1-Luc/VSV-G (MOI = 2) or mock infected. NVP was used to block HIV-1 reverse transcription. THP-1 cells were harvested at 1-4 dpi and differentiated THP-1 cells were harvested at 1-2 dpi for analyses. ROS was measured by H2DCFDA staining and flow cytometry. Unst., Unstained cells.

## Notes

### Competing Interest Statement

The authors have declared no competing interest.

